# Spatial growth in food-like matrices differentially modulates food-related stress responses but enhances digestive tolerance in major foodborne pathogens

**DOI:** 10.64898/2026.03.05.710002

**Authors:** Elodie Hoch, Christina Nielsen-Leroux, Laurent Guillier, Bernard Hezard, Romain Briandet, Lysiane Omhover-Fougy

## Abstract

Foods are spatially structured and heterogeneous matrices in which microbial pathogens predominantly grow as immobilised microcolonies rather than planktonic free cells. However, most predictive microbiology and risk assessment models rely on homogeneous liquid cultures, potentially overlooking spatial effects on stress adaptation. Here, we investigated how growth within food-like semi-solid matrices influences stress adaptation and digestive tolerance of major foodborne pathogens. We compared planktonic and spatialised lifestyles across multiple species exposed to salt and organic acid stresses. Spatial growth profoundly altered growth dynamics in a stress- and species-dependent manner. Notably, spatial growth markedly enhanced tolerance to simulated gastrointestinal stresses *in vitro*, particularly under acidic conditions. This protective effect was further confirmed *in vivo* within the acidic midgut of *Hermetia illucens* larvae. Our findings demonstrate that spatial organisation generates distinct physiological states that increase pathogen resilience, highlighting the need to integrate spatialisation into predictive models and quantitative microbial risk assessment.

## 1. INTRODUCTION

Food products are one of the main vehicles for the transmission of pathogenic microorganisms to humans. Despite substantial improvements in hygiene, food processing and risk management, foodborne diseases remain a significant public health burden worldwide (www.who.int). In the European Union alone, 6,558 foodborne outbreaks were reported in 2024, representing a 14.5% increase compared to 2023. Campylobacteriosis, salmonellosis, STEC (Shiga toxin *Escherichia coli*) infection and listeriosis remain the most frequently reported, with Salmonella enterica being the leading causative agent in multi-country outbreaks^1^. Although less frequent, *Listeria monocytogenes* infections are associated with the highest hospitalisations and fatality rates, reaching 30% in invasive forms and often leading to severe neurological sequelae^2^. Before reaching the human host, foodborne pathogens are exposed to a succession of environmental constraints along the food chain, including food processing, formulation, storage and distribution^3^. These environments subject bacteria to multiple stresses such as disinfectants, heat treatments, reduced water activity, organic acids, salts and other preservation hurdles. In addition, pathogens must subsequently withstand the physicochemical constraints of the gastrointestinal tract, including low pH, bile salts and oxidative stress. The ability of foodborne bacteria to adapt to these stresses is therefore a critical determinant of both food safety and infection risk.

Moreover, foods are not homogeneous environments. They encompass a wide range of physical states, from liquids to semi-solid and solid matrices, including gels, emulsions and multiphasic systems (e.g., mayonnaise, cream)^4^. These structures generate gradients of nutrients, pH, oxygen and water activity that evolve over time and in response to microbial activity. Experimental studies^5^ have shown that the physical structure and the presence of heterogeneous microenvironments on food matrices strongly influence bacterial spatial organisation and promote phenotypic diversification^4,6^. Consequently, microorganisms growing in foods are often immobilised and develop as single cells, small aggregates or embedded microcolonies rather than as freely suspended planktonic populations ^7^.

Nevertheless, most studies investigating stress adaptation in foodborne pathogens have relied on planktonic liquid cultures, which fail to capture microscale heterogeneity and physical constraints characteristic of real foods^8–10^. Consequently, the physiological states generated by spatial confinement within food matrices remain largely underexplored, creating a critical gap between experimental models and real food ecosystems. This methodological gap limits our ability to accurately predict bacterial behaviour in structured food systems. The impact of spatial structure on microbial physiology has been extensively investigated in biofilms and colony-biofilm models. In these systems, spatial organisation generates steep gradients of nutrients, oxygen and metabolic by-products, leading to stratified physiological states and the emergence of stress-tolerant subpopulations. Biofilm-associated bacteria display enhanced tolerance to antimicrobials, disinfectants and environmental stresses, partly due to the presence of extracellular polymeric substances, reduced growth rates, metabolic heterogeneity and collective stress responses. These responses have been shown in both submerged biofilm models^11–18^ and in colony-biofilm models^19–22^.

While these models highlight the impact of spatial structure on bacterial behaviour, they differ fundamentally from the embedded microcolonies that develop within food matrices. Food-like semi-solid models have begun to bridge this gap^23,24^. However, systematic comparisons across multiple major pathogens and a diversity of food-related stresses remain scarce, particularly when integrating both growth dynamics and post-ingestion tolerance. Boons et al. 2014, showed that in hydrogel matrices containing both gelatine and dextran phases (a glucose polymer), *E. coli* preferentially develops as colonies within the dextran phase, regardless of which phase is dispersed^25^. Saint Martin et al., 2023 demonstrated that microcolonies of *E. coli* O157:H7 cultivated in hydrogels were not affected by a 4-hour exposure to HCl at pH 2.0 (mimicking gastric acid stress), whereas their planktonic counterparts were eradicated after this treatment^26^. A similar behaviour has been demonstrated in probiotic bacteria organised into hydrogel microcolonies compared to their planktonic counterparts^27^.

Beyond its impact on growth dynamics, the spatial mode of bacterial growth can also influence virulence^17,28,29^. Indeed, the physiological state of bacterial cells is closely linked to the expression of virulence factors, which are often regulated by environmental conditions such as nutrient availability, stress exposure, and cell density. In structured environments, such as biofilms or microcolonies, spatial heterogeneity generates subpopulations with distinct metabolic states, including slow-growing or dormant cells that may exhibit increased stress tolerance and altered virulence potential^3,30–32^. These observations suggest that bacterial organisation within food matrices may not only affect survival under stress but also modulate the virulence of ingested pathogens.

In this context, the present study investigates how spatial growth within food-like matrices, affects stress adaptation and digestive tolerance in major foodborne bacterial pathogens. Using a high-throughput experimental framework, we compared planktonic and spatialised lifestyles across a diverse panel of pathogens exposed to food preservation stresses, focusing on salt and organic acids. We further assessed whether spatialised growth promotes tolerance to gastrointestinal stresses using complementary *in vitro* digestion assays and an *in vivo* insect larval model. We hypothesise that spatialised microcolony growth generates unique physiological states that enhance bacterial resilience compared with homogeneous planktonic cultures. By addressing this hypothesis, this work aims to improve our understanding of pathogen behaviour in structured foods and to highlight the importance of spatial organisation for food safety risk assessment.

## 2. RESULTS

### 2.1. Spatial structure alters bacterial growth responses to food-related stresses

Quantitative growth parameters were used to evaluate the impact of spatial organisation on bacterial responses to food-related stresses. Lag phase duration (Lag) and maximum growth rate (*µ*_max_) were extracted using the Baranyi and Robert’s model for both planktonic and spatialised lifestyles (see **Supplementary material 1**). While planktonic growth was monitored by homogeneous turbidimetry, spatialised growth in hydrogels was characterized by multipoint optical density measurements, allowing the integration of spatial heterogeneity into growth parameters estimation.

To explore global patterns across all bacteria-stress combinations, a non-supervised K-means clustering analysis was performed using Lag and *µ*_max_ values from all experimental conditions, with 9 replicates per condition (**Figure 1A**). Based on the within-cluster sum of squares (WSS) criterion, three clusters were identified as the optimal partition of the dataset. Cluster 1 was characterised by a long Lag phase (25 h to 60 h) associated with low *µ*_max_ values (<0.1 h^-1^). In contrast, cluster 3 groups conditions with high *µ*_max_ values (0.2 h^-1^ to 0.6 h^-1^) and a smaller Lag phase (<20 h). Cluster 2 displays intermediate growth behaviour, with Lag phases shorter than 20h and *µ*_max_ values below 0.2 h^-1^.

To assess the contribution of experimental variables to cluster structure, the distributions of strains, stresses, and lifestyles within each cluster were quantified (see **Supplementary material 2**). Among these variables, lifestyle emerged as the dominant factor discriminating between the clusters (**Figure 1B**). Planktonic conditions were strongly overrepresented in cluster 3 (97 %) and underrepresented in clusters 1 (17%) and 2 (38%). Conversely, spatialised conditions predominated in clusters 1 (83%) and 2 (62%), whereas they were almost absent in cluster 3 (3%).

This distribution indicates that spatialised growth is primarily associated with growth profiles characterised by extended Lag phases and reduced or intermediate growth rates, whereas planktonic growth is associated with rapid growth dynamics. Overall, these results demonstrate that spatial structure is a major determinant of bacterial growth responses under food-related stress conditions. Importantly, this global segregation of growth profiles according to lifestyle was observed across phylogenetically diverse pathogens, suggesting that spatialisation acts as a unifying ecological driver beyond species-specific physiology.

**Figure 1:**
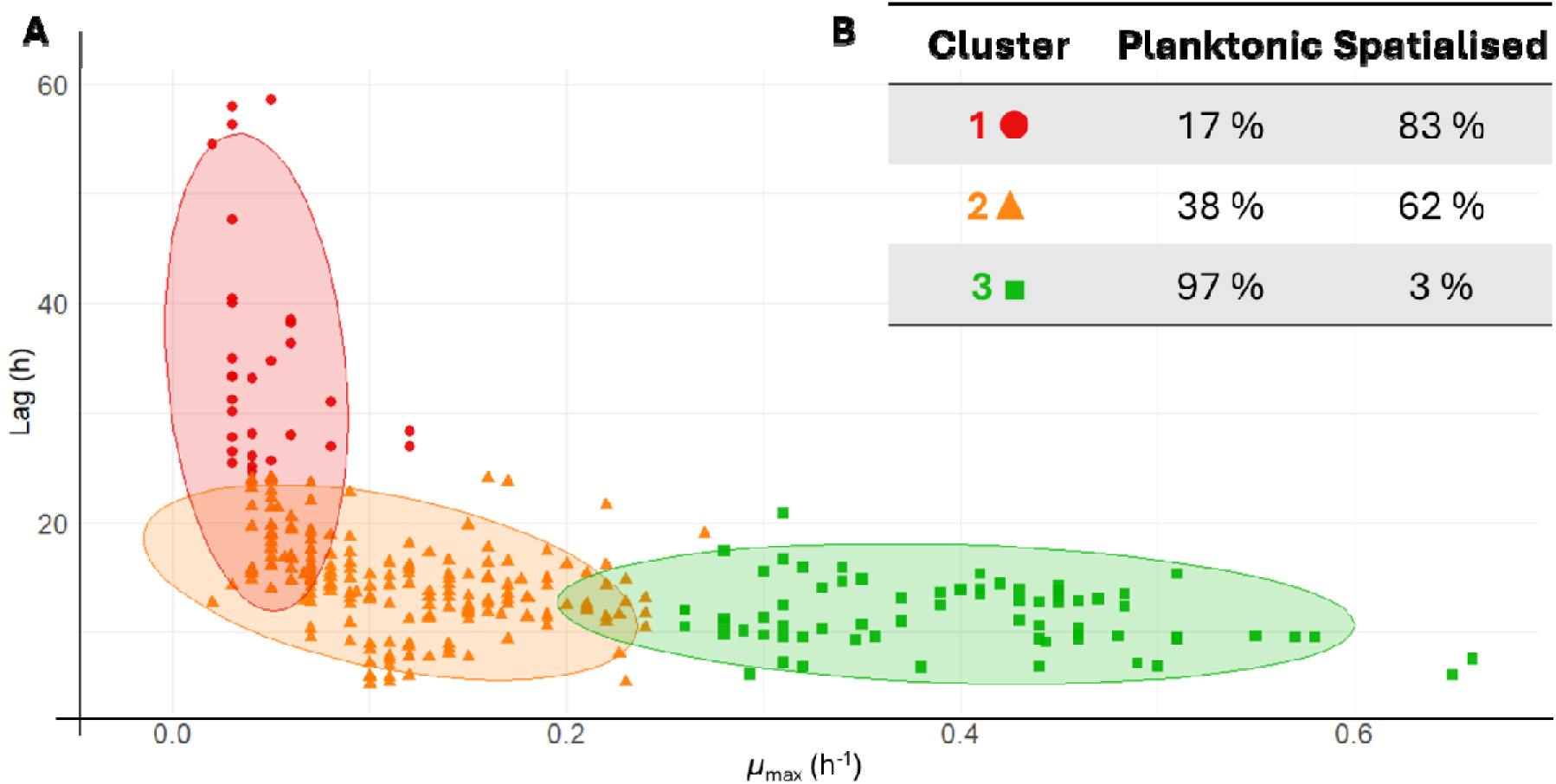
**A**: K-means clustering of microbial growth parameters (Lag and *µ*_max_) under various environmental stresses. Each point represents a bacterial strain grown under specific conditions (a combination of food stress and lifestyle), with values corresponding to the mean of biological replicates (n = 9), for two quantitative parameters: Lag (duration of the Lag phase) and *µ*_max_ (maximum growth rate). The optimal number of clusters (k = 3) was selected using the elbow method based on within-cluster sum of squares (WSS). Red circles ● correspond to cluster 1; orange triangles ▲ correspond to cluster 2, and green squares ▪ correspond to cluster 3. **1B**: Table showing the distribution of lifestyle across the 3 clusters.

### 2.2. The protective effect of spatialisation is highly dependent on the bacteria-stress combination

To specifically evaluate the effect of spatial organisation on bacterial sensitivity to food-related stresses, growth responses were analysed using ΔLag and Δ*µ*_max_, defined as the difference between growth parameters measured under control and stress conditions. For each bacteria-stress pair, the impact of spatialisation was quantified by calculating the difference between spatialised and planktonic responses ΔLag_spatialised_ - ΔLag_planktonic_ and Δ*µ*_maxspatialised_ - Δ*µ*_maxplanktonic._ These differences are represented in **Figure 2**.

In these representations, red values indicate increased sensitivity to stress under spatialised conditions, reflected by a longer Lag phase or a reduced maximum growth rate compared to planktonic cultures. Conversely, green values indicate a protective effect of spatialisation, characterised by a reduction in Lag phase duration and/or an increase in *µ*_max_.

The impact of spatialisation on growth parameters is markedly dependent on the bacterial species, the type of stress and its intensity. For several bacteria-stress combinations, responses were strongly concentration-dependent. This was particularly evident for Se LT2, for which spatialised growth conferred significant protection at 6.00%, while an opposite effect was observed at 2.00% NaCl. At intermediate salt concentration (3.00% and 4.50% NaCl), no significant effect of spatialisation on the Lag phase duration was detected.

A similar concentration-dependent response was observed for Lm EGDe, which exhibited a significant reduction in sensitivity to salt stress at 3.00%, 4.50% and 6.00% NaCl, whereas no significant effect was observed at 2.00% NaCl. For this strain, the magnitude of the spatialisation effect increased with salt concentrations, with the strongest responses observed at 4.50% and 6.00%. In contrast, some bacteria displayed divergent responses depending on the growth parameter considered. For Bc and Vp exposed to increasing concentrations of acetic acid, spatialised growth was associated with an increase in *µ*_max,_ accompanied by a concomitant extension of the Lag phase.

Overall, the most pronounced protective effects of spatialisation on growth rate were observed under salt stress conditions (**Figure 2B**), particularly at 6.00% NaCl. In comparison, the effect of specialisation on Lag phase duration (**Figure 2A**) was more variable and strongly dependent on the specific bacteria-stress combination. Based on these observations, salt stress showed the strongest spatialised protection responses. Moreover, the presence of salt does not affect the hydrogel texture, that is why it was selected for further experiments.

For subsequent experiments, representative bacteria-stress combinations were selected based on the magnitude and diversity of responses. Lm EGDe and Ye were selected due to their strong protective response to 6.00% NaCl. For Ye and Se LT2, both low (2.00%) and high (6.00%) salt concentrations were retained to capture threshold effects. Bc was also included at 6.00% NaCl due to its contrasting responses on Lag phase and growth rate. Vp also showed interesting responses to salt stress, but known to be halophilic, it was not included in further investigation.

**Figure 2:**
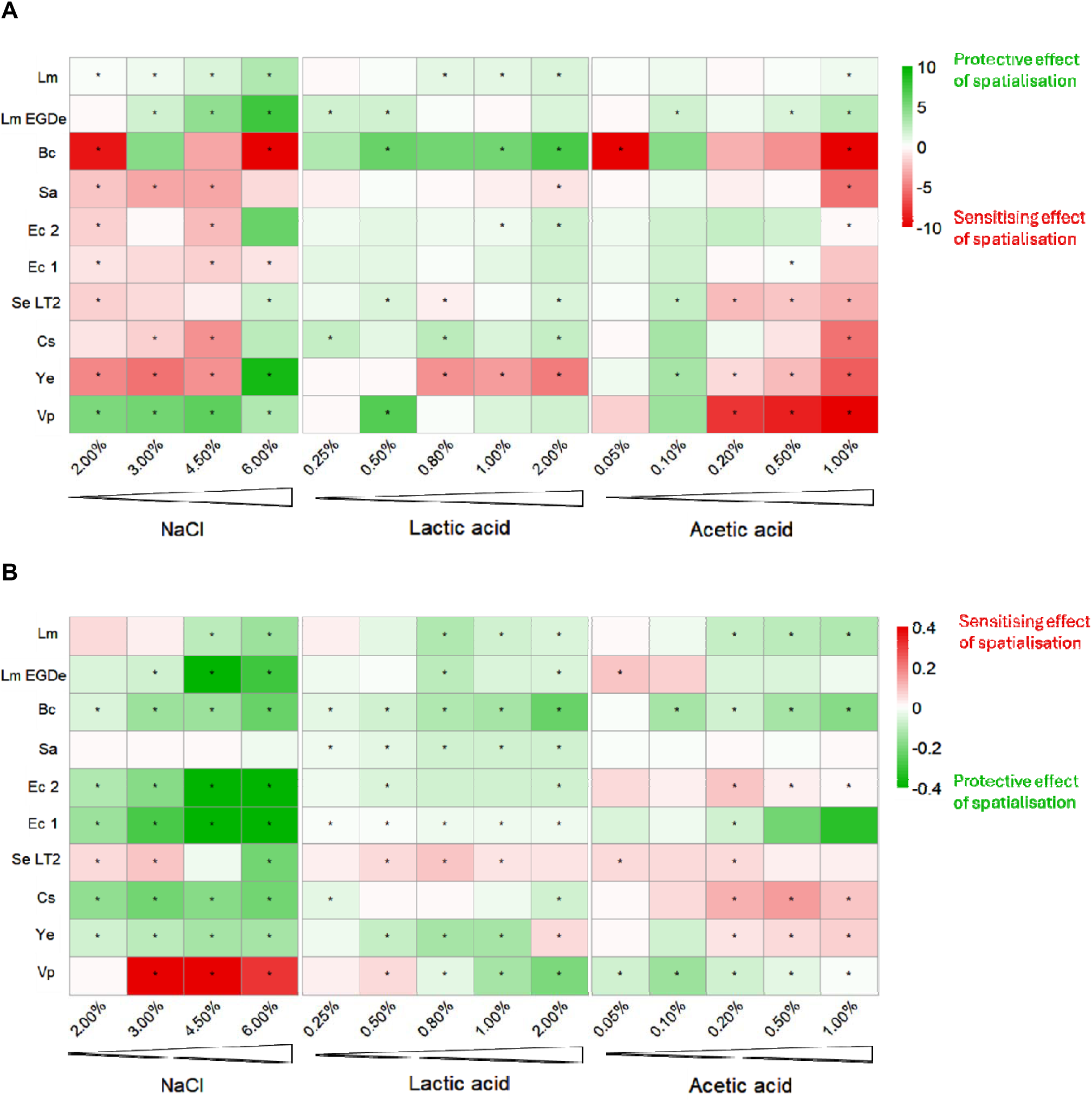
Influence of spatial structure on Lag phase (**A**) and *µ*_max_ (**B**) responses of foodborne bacterial pathogens under food-related stress (n = 9). Bacterium-stress pairs are presented with different levels of food stress (NaCl, lactic acid, and acetic acid) on the x-axis, and various pathogenic strains on the y-axis. Scales correspond to the differences between spatialised delta and planktonic delta, where delta represents the difference between growth data under controlled conditions and under stressed conditions. Red values indicate greater food stress sensitivity in a spatialised environment, while green values reflect a protective effect of spatialisation. Growth parameters are extracted from Baranyi’s model. Pairwise comparison using a parametric or non-parametric test was applied between lifestyles. Asterisks indicate significant differences between lifestyles (p<0.05).

### 2.3. Spatialised growth in food-like matrices leads to the formation of pathogen-specific microcolonies architecture that are weakly affected by salt stress

Bacterial pathogens studied here are characterised by morphological and metabolic pathways diversity^44^. To demonstrate phenotypic diversity in hydrogels, four strains and two media were selected. Macroscopic images acquired using a robotised imaging platform (see **Supplementary Material 3**) and to characterise microcolony architecture at a finer spatial scale, three-dimensional observations were performed using confocal laser scanning microscopy (CLSM). Microcolonies were stained with the cell-permeant nucleic acid dye Syto 61, allowing visualisation of their internal structure and morphology (**Figure 3A**).

During growth, all tested pathogens formed microcolonies within the bulk of the GSFM and S-GFSM. These structures, hereafter referred to as microcolonies, consist of clusters of cells spatially confined within the hydrogel, and are distinct from freely suspended populations. The size and morphology of microcolonies varied according to the bacterial species, the composition of the hydrogel, and the presence of environmental stresses such as elevated NaCl concentrations or organic acids, as well as the initial inoculum density. In addition, large colonies were occasionally observed at the air-gel interface, corresponding to macrocolonies developing under conditions of increased oxygen availability. The relative proportion of microcolonies and macrocolonies differed between species. Lm EGDe predominantly formed small and regularly shaped microcolonies, whereas Bc frequently produced larger colonies at the gel surface. The composition of the hydrogel also influences the preferred growth organisation. For Ye, microcolony formation was more prominent in S-GSFM, whereas Se LT2 preferentially formed microcolonies in GSFM. In addition, Se LT2 exhibited the formation of gas bubbles within the hydrogel, consistent with the fermentative metabolism characteristic of Enterobacteriaceae. Within the observed field, microcolonies were randomly distributed and did not display preferential spatial location within the hydrogel. When hydrogel was supplemented with 6.00% NaCl, the number of detectable microcolonies was reduced compared to the control condition (GSFM), while their spatial distribution remained homogeneous. Among the strains analysed, Bc exhibited noticeably larger microcolonies than the other pathogens. In several instances, the size of these structures exceeds the optical penetration depth of the laser, resulting in partial visualisation of the microcolony interior.

Quantitative analysis of microcolonies’ architecture was performed by extracting geometric parameters, including biovolume and sphericity, from segmented CLSM images (**Figure 3B**). For all strains, the mean biovolume of microcolonies ranges between 4 and 5 log_10_ µm^3^, while mean sphericity values ranged from 0.7 to 0.9 (A.U.). The presence of 6.00% NaCl in the hydrogel did not significantly modify the biovolume and the sphericity of microcolonies for Se, Lm, or Ye. In contrast, Bc exhibited a significant increase in microcolonies biovolume under saline conditions, with a mean value increasing from 4.06 ± 0.19 log_10_ µm^3^ to 4.68 ± 0.62 log_10_ µm^3^. This increase was associated with a shift towards a more lenticular morphology, reflected by a significant decrease in sphericity from 0.83 ± 0.05 to 0.70 ± 0.14 (A.U.).

**Figure 3:**
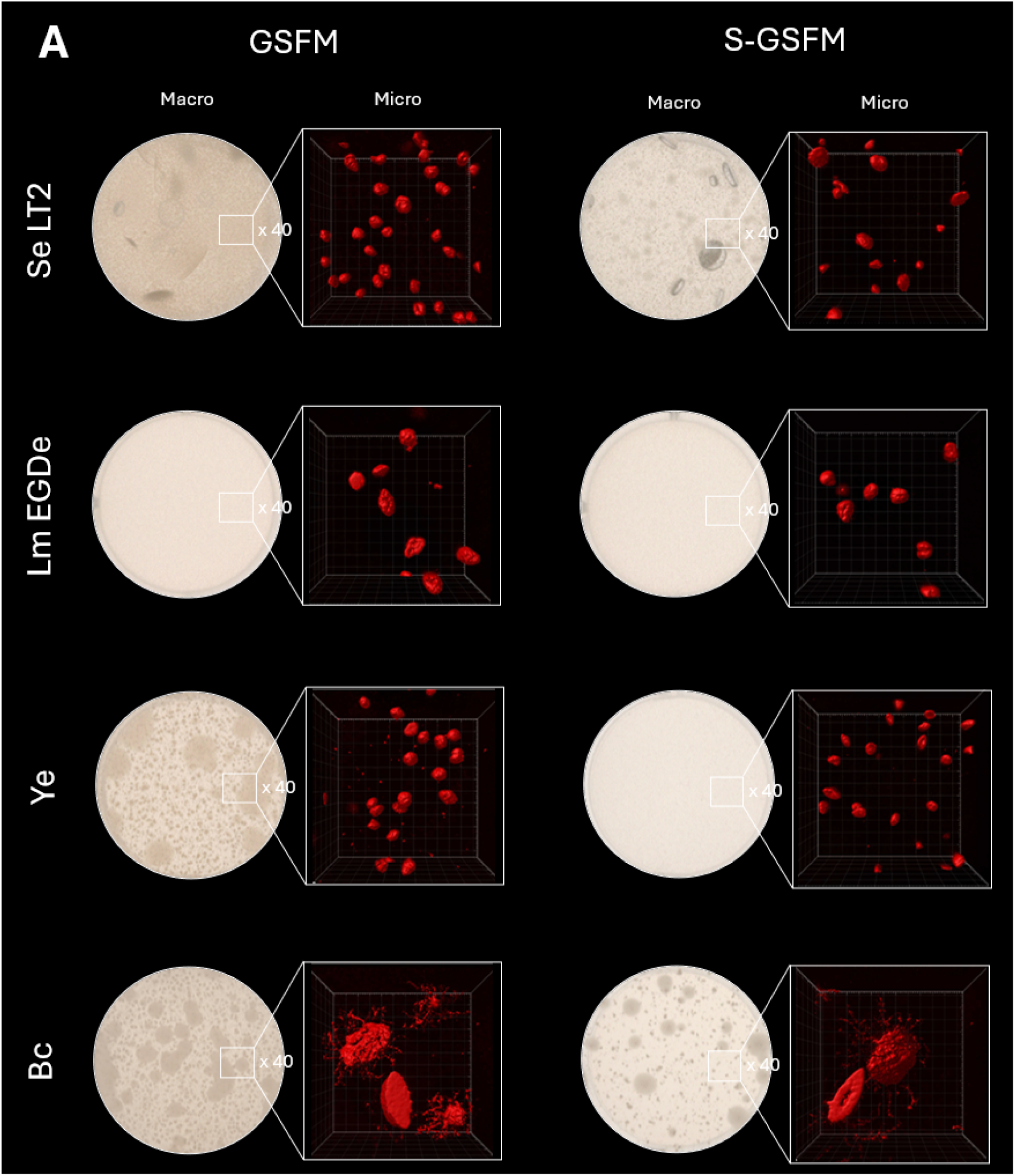
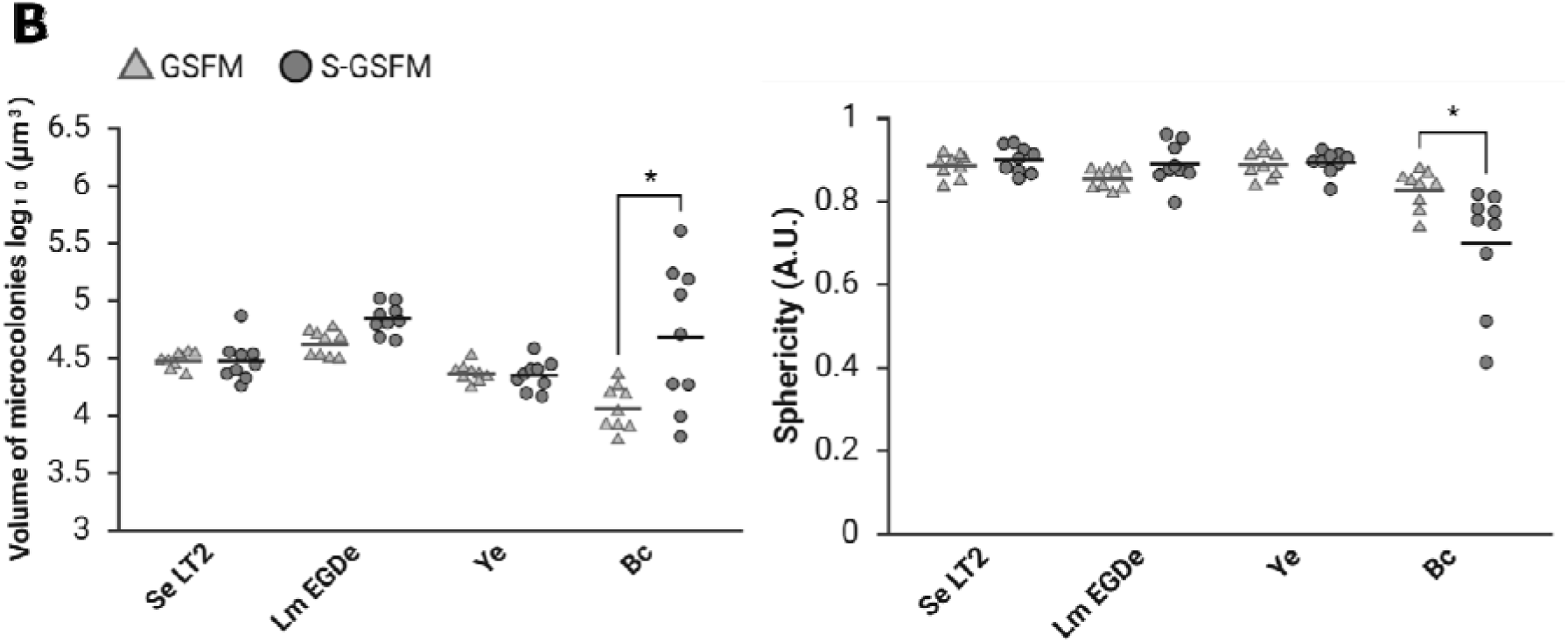
Microcolonies architecture in GSFM (▲) or S-GSFM (●), after 96h at 25°C. **A**: Macroscopic images are obtained using an automated robot that integrates incubation and high-resolution imaging. Microscopic observations of microcolonies, endpoint stained with the cell-permeant nucleic acid dye Syto 61, are obtained with confocal laser scanning microscopy. Representative pictures correspond to the microcolonies isosurface after fluorescence segmentation using the Imaris isosurface functions. **B**: Quantitative parameters extracted from the isosurface of microcolonies in Imaris software. Each point represents an independent replicate (n=9), bars correspond to the mean of volume of microcolonies log_10_ (µm^3^) and sphericity (A.U.). A two-way ANOVA was performed to assess the effect of food stress. When a significant effect of the environment was detected, the Bonferroni multiple comparisons test was applied to evaluate the level of significance. Asterisks indicate significant differences between environments (p<0.05).

### 2.4. Spatialised growth enhances tolerance to simulated digestive stresses *in vitro*

To determine whether spatialised growth influences bacterial tolerance to conditions mimicking the gastrointestinal tract, four bacterial pathogens (Se LT2, Ye, Lm EGDe, and Bc) were exposed to *in vitro* digestive stresses following growth under planktonic or spatialised conditions. Bacteria were harvested after four days of incubation in the stationary phase and subsequently subjected to HCl stress (pH 2), oxidative stress (10 mM H_2_O_2_), or bile salts (1%) (**Figure 4**).

For all strains, exposure to acidic conditions resulted in the largest reduction in cultivability in planktonic cultures. In contrast, growth as microcolonies markedly increased survival under HCl stress for Se LT2 and Ye, with log reduction decreasing from approximately 5-7 log_10_ CFU/mL in planktonic cultures to less than 1 log_10_ CFU/mL under spatialised conditions (p < 0.05) (**Figures 4a and 4b**). For Lm EGDe and Bc, a protective effect of spatialisation against acidic stress was observed only when bacteria were previously grown in S-GSFM, whereas no significant difference between lifestyles was detected in the control medium (**Figures 4c and 4d**).

Responses to bile salts were more heterogeneous. No significant effect of spatialisation was observed for Lm and Bc when bacteria were grown in the control medium. However, when previously exposed to saline conditions, these strains exhibited increased sensitivity to bile salts in the spatialised state. In contrast, Se LT2 displayed a significant protective effect of microcolonies growth against bile salts following growth in S-GSFM.

Oxidative stress responses also varied between species and growth conditions. Se and Bc were consistently more tolerant to H_2_O_2_ when grown as microcolonies, irrespective of the growth medium. In contrast, Ye and Lm exhibited protective effects of spatialisation only under specific growth conditions (S-GSFM for Ye and GSFM for Lm).

Overall, among the four strains tested, Se LT2 and Ye ATCC 27729 exhibited the highest tolerance to simulated digestive stress when grown as microcolonies, with log reduction close to 1 log_10_ CFU/mL across conditions. Se LT2 showed the largest difference between lifestyles, particularly under HCl stress (∼7 log differences), followed by oxidative stress (∼3-4 log) and bile salts (∼1.5 log).

**Figure 4:**
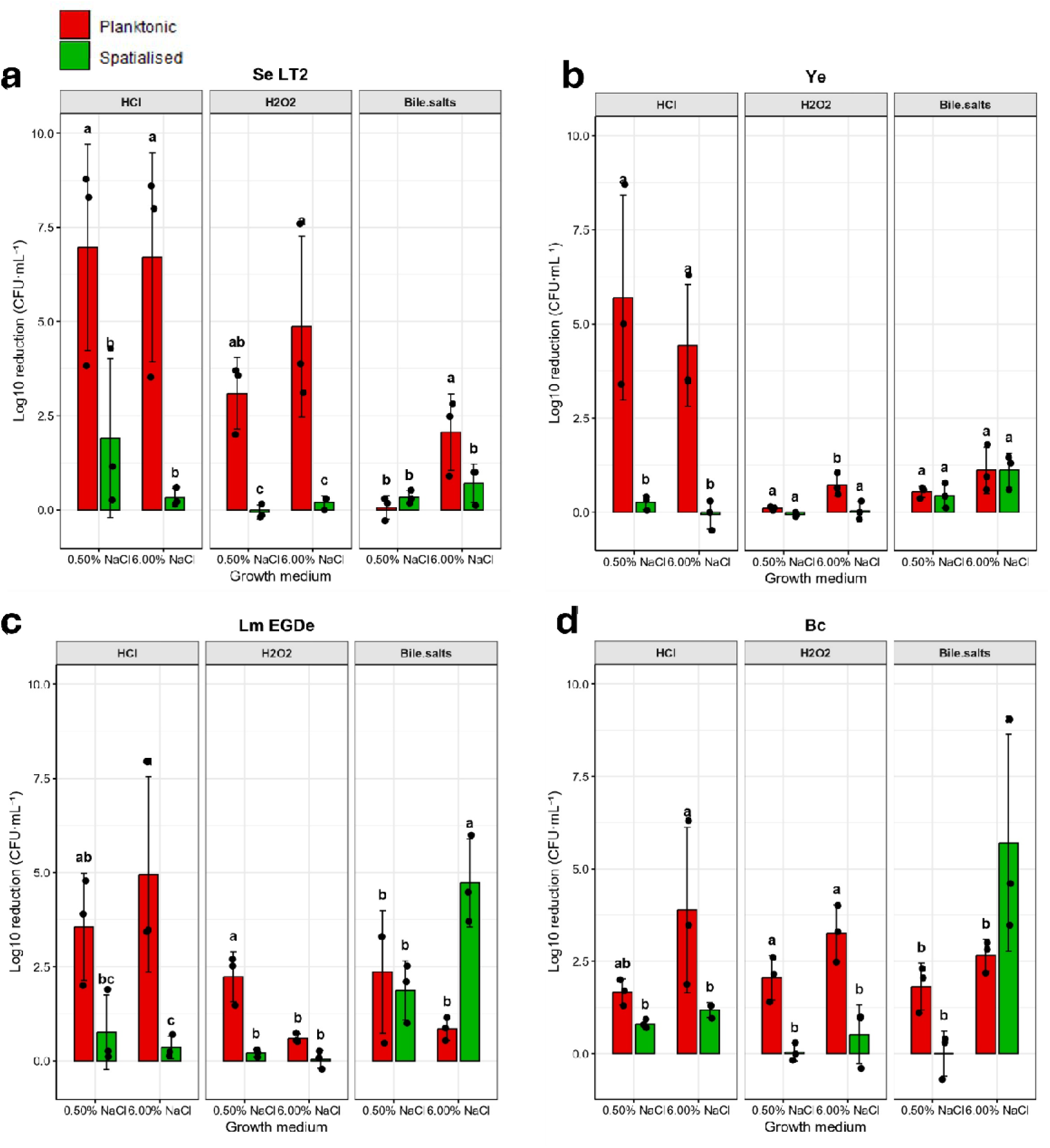
Survival of planktonic cultures (red) and hydrogel microcolonies (green) of (**a**) Se LT2, (**b**) Ye, (**c**) Lm EGDe, and (**d**) Bc, upon digestive stress (HCl: pH 2.0; hydrogen peroxide H_2_O_2_: 10 mM; bile salts: 1%). The log reduction after treatment was calculated by subtracting the bacterial concentration in the unstressed condition from that in the digestive stress. Statistical analysis consisted of two-way ANOVA. When a significant effect of lifestyle was detected, Fisher’s Least Significant Difference (LSD) post hoc test was applied to determine which digestive and food stressors. Stress groups with different alphabets indicate a significant difference (p < 0.05).

### 2.5. *In vivo* ingestion reveals enhanced survival of spatialised Salmonella

Based on the *in vitro* digestive stress results, an *in vivo* experiment using the same environments, planktonic and microcolony lifestyles, as tested previously, was conducted with the Se LT2 strain. The larvae of *H. illucens* are described as having a digestive tract in which the pH varies widely across regions ^43^. The larval gut is composed of three parts: foregut, midgut and hindgut. The digestion and absorption take place in the midgut region. The midgut region is divided into three parts having pronounced pH shifts: the anterior part has a pH around 7, the middle part has a pH less than 3 and the pH increases again toward pH 8 at the posterior part (see **Supplementary Material 4**). The focus here is therefore on the transition to pH 2 in the midgut. The intestinal pH was measured using pH paper to verify the presence of an acidic region (see **Supplementary Material 5**).

In a first experiment used as a control, after 24 hours of ad libitum ingestion (see **Supplementary Material 6**), larvae were collected and crushed to release any *Salmonella* present in the gut. The presence of *Salmonella* in/on the hydrogel promotes ingestion by *H. illucens* but the bacterial lifestyle does not change the time of total ingestion. The microbiological counts are presented in **Table 3**. No *Salmonella* were enumerated at T0 and T24 in *H. illucens.* The initial Se LT2 count in the hydrogel was approximately 10 log_10_ CFU for all conditions tested. Without larvae, the *Salmonella* spp. count remained unchanged (data not shown), indicating that neither growth nor death occurred during the ingestion period. After ingestion, *Salmonella* spp. counts in the substrate were respectively 7.83 ± 0.61 log_10_ CFU and 8.27 ± log_10_ CFU for the planktonic and spatialised environment. With 6.00% NaCl supplementation, *Salmonella* found in the hydrogel after larval consumption were 8.08 ± 0.36 log_10_ CFU in planktonic and 8.23 ± 0.37 log_10_ CFU in spatialised environment. These data indicated that *H. illucens* might have ingested the same quantity of *Salmonella* regardless of the lifestyle or the medium composition. However, it is also possible that the enumerated pathogen, passed through the digestive tract, survived, and was excreted into the Petri dish and mixed up, mixing with the non-ingested bacteria.

Focusing on the whole non-dissected larvae, no significant differences in *Salmonella* levels were detected between the various conditions. Approximately 8.0 - 8.5 log_10_ CFU were enumerated inside larvae, distributed throughout all portions of the digestive tract. No significant differences between the two lifestyles and the two media tested were observed in the concentration of Salmonella detected in *H. illucens*. Nevertheless, we observed a *Salmonella* total count evolution to 10 log_10_ CFU at T0, which was higher than at T24, where a *Salmonella* count of 8.0 - 8.5 log_10_ CFU (sum of hydrogel and larvae count) was reached. This high number of *Salmonella* found indicates that the larvae’s intestines provide a suitable niche for the pathogen to colonise *H. illucens*. Yet the difference observed between T0 and T24 in the global pathogen count of interest is presented in the last column of **Table 3**. These numbers indicate a decrease in the number after ingestion, indicating a *Salmonella* death.

This first *in vivo* experiment demonstrates that *H. illucens* can serve as an *in vivo* model for ad libitum ingestion of planktonic cells and hydrogel microcolonies of *Salmonella*. Because no significant differences were detected in the entire gut of *H. illucens*, we decided to focus on the acidic region of *Hermetia’s* midgut (see **Supplementary Material 4**).

The Se LT2 load was enumerated after dissection, within the midgut at pH 3 using a chromogenic medium (**Figure 5**). The counts show that the load in *Salmonella* is very heterogeneous within the same condition. In planktonic or spatialised conditions, there are no significant differences; the average load is 3 log_10_ CFU/larva. In planktonic 6.00% NaCl, the average load is 1.8 log_10_ CFU/larva, whereas in spatialised lifestyle, it is 3.5 log_10_ CFU/larva. In the presence of 6.00% NaCl in microcolonies, Se LT2 are significantly more tolerant to the midgut of *H. illucens,* compared to the free cells form. As demonstrated in **Table 1**, the global charge in Se LT2 in the gut did not differ according to the lifestyle. In the second experiment (**Figure 5**), the dissection of the midgut, we show the same protective effect of spatialisation against acidic pH as in the *in vitro* model.

**Table 1:**
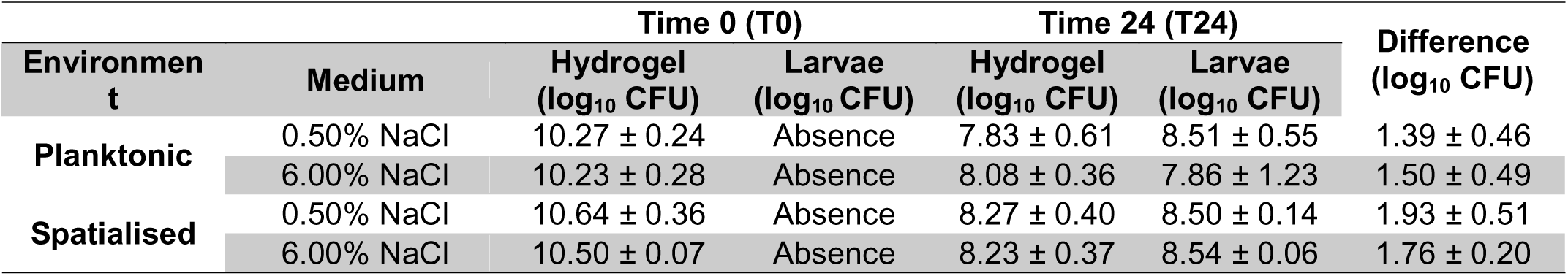
Total *Salmonella* spp. counts from *H. illucens* whole larva and hydrogel. All conditions were incubated separately at 25°C. Values represent the mean (± standard deviation) of four batches, each with 3 larvae per condition (n=4). Values are expressed as log_10_ CFU per Petri dish (= condition).

**Figure 5:**
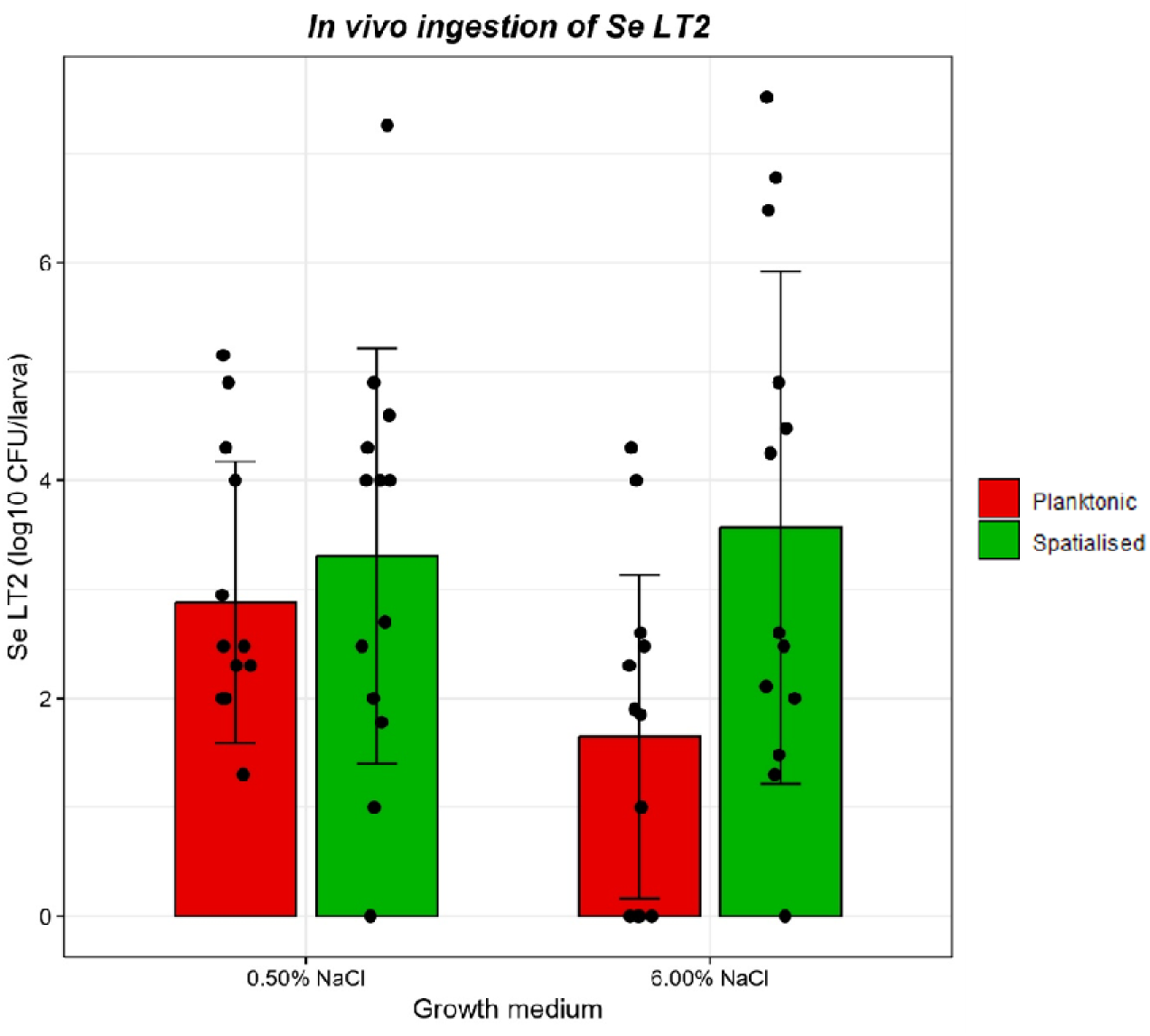
Survival in the acidic gut region, of Se LT2 planktonic cultures (red) and hydrogel microcolonies (green) in medium containing 0.50% or 6.00% NaCl, in the acidic region of the gut of *H. illucens* after 24h ad libitum ingestion. Statistical analysis consisted of the Mann-Whitney test to determine which conditions are significantly different. Stress groups with different alphabets indicate a significant difference (p<0.05).

Beyond descriptive differences in growth parameters, our data demonstrate that the spatial history of pathogens before ingestion significantly influences their survival under gastrointestinal constraints. This observation has important implications for predictive microbiology and quantitative microbial risk assessment frameworks.

## 3. DISCUSSION

Understanding how food microstructures shape the physiological state of foodborne pathogens is critical for accurately predicting their survival during food conservation and their passage through the gastrointestinal tract. In this study, we demonstrate that spatialised growth within food-like semi-solid matrices profoundly modifies bacterial stress response and digestive tolerance compared with conventional planktonic cultures. By combining high-throughput growth analysis with *in vitro* and *in vivo* digestive stress models, we show that spatial organisation acts as a key driver of bacterial resilience, with effects that depend strongly on both the pathogen and the nature of the applied stress.

Growth in agarose-based food-like matrices led all tested pathogens to form immobilised microcolonies rather than as dispersed single cells. Macroscopic observations revealed heterogeneous microcolonies embedded within the hydrogel, with occasional larger colonies at the air-gel interface, highlighting the strong influence of spatial constraints imposed by the semi-solid matrix. At the microscopic scale, confocal imaging revealed that microcolony morphology and size varied with bacterial species and stress conditions. Under control conditions, most strains formed compact, ovoid to quasi-spherical microcolonies with relatively homogeneous spatial distribution. Exposure to saline stress induced strain-dependent architectural changes, most notably in Bc, which formed larger, more lenticular microcolonies at high NaCl concentrations. In contrast, Se LT2, Lm, and Ye maintained comparable morphologies across conditions. This diversity is most likely due to different regulatory mechanisms ^45–47^. Moreover, the lenticular shape induces thickness differences within microcolonies, possibly with nutrient and metabolic gradients, leading to phenotypic diversification of embedded bacteria between the spherical and lenticular shapes.

A central outcome of this study is that spatialised growth markedly alters bacterial responses to food-related stresses, but in a manner that depends on both stress intensity and bacterial species. The Baranyi and Roberts model was used for its ability to model the Lag phase and the transition to the exponential phase in isothermal conditions. Using growth parameters as quantitative indicators of stress adaptation, we show that salt stress provides the clearest illustration of the protective effect of spatial organisation. For several pathogens, including Se LT2 and Lm EGDe, spatialised growth significantly reduced Lag phase duration and increased maximum growth rate at high NaCl concentrations, whereas little or no effect was observed at lower salt levels. This concentration-dependent response suggests that spatial structuring confers a selective advantage primarily under severe osmotic stress^48^. One plausible explanation is that diffusion limitations within microcolonies attenuate the effective stress experienced by interior cells, generating microenvironments that allow partial growth and adaptive responses even when external conditions are harsh^24,49,50^. Similar mechanisms have been described in biofilms and macrocolonies, where gradients of nutrients, oxygen, and stressors lead to the coexistence of subpopulations with distinct metabolic states^20,51,52^. Our results extend this concept to embedded microcolonies in food-like matrices. In contrast, responses to organic acid stresses were more variable and, in some cases, contradictory between growth parameters. As previously reported, changes in gel texture and porosity can affect diffusion and bacterial immobilisation, making it difficult to disentangle purely physiological effects from matrix-mediated ones. These findings underline the importance of carefully considering matrix–stressor interactions when interpreting microbial behaviour in structured foods.

It is important to note that the improved stress tolerance observed in a spatialised culture is probably multifactorial. As mentioned before, diffusion limitations phenomena may reduce the effective intensity of external stresses inside microcolonies, allowing cells to experience exposure that enhances adaptive responses^24,50^. Another point is that the growth rate heterogeneity induced by spatialisation can promote slow-growing subpopulations, which are intrinsically more tolerant to a wide range of stresses^5,53,54^. Finally, localised nutrient limitation and redox gradients may activate global stress responses (RpoS in *Salmonella* and *E. coli*), leading to cross-protection against unrelated stresses. Such mechanisms have been described in macrocolonies and biofilms as collective responses that enhance robustness biofilms^55–57^.

Beyond food preservation stresses, a major outcome of this work is the demonstration that spatialised growth before ingestion enhances bacterial survival to simulated gastrointestinal conditions (**Figure 6**). *In vitro* digestion assays revealed a strong protective effect of the microcolony lifestyle against acidic stress, particularly in Se LT2 and Ye, for which planktonic populations were almost completely inactivated, whereas spatialised populations retained substantial cultivability. Acid tolerance is a critical determinant of pathogen survival during gastric passage, and our results suggest that spatial organisation induces pre-adaptive physiological states that enhance resistance to low pH ^58^. This pre-adaptation likely emerges from the combined effects of diffusion-limited stress exposure, metabolic stratification, and the enrichment of slow-growing subpopulations within microcolonies. This phenomenon may involve partial activation of acid resistance systems, global stress regulators such as RpoS, or the presence of slow-growing or metabolically altered subpopulations within microcolonies. Similar pre-adaptation effects have been described in biofilms and colony biofilms, where spatial heterogeneity promotes cross-protection against multiple stresses^27,50,59^.

Responses to oxidative stress reveal a more nuanced picture; it is likely governed by species-specific regulatory mechanisms, and spatial growth does not uniformly enhance defences against reactive oxygen species (ROS). Moreover, as mentioned before, osmotic and oxidative stresses encountered in foods can activate global stress responses, thereby increasing tolerance to gastric acidity and bile salts. This pre-adaptation phenomenon is known in *L. monocytogenes*^60^ and is correlated with our results in the planktonic lifestyle. A hypothesis done by Begley, Gahan, and Hill 2005, is that the presence of microenvironments created by the food matrix in the intestine can affect the bacteria’s survival by binding bile acids to food constituents and therefore preventing toxicity^61^. Our findings are correlated with this hypothesis, because we show a reduction in the effect of bile salts in embedded microcolonies of Se LT2 and Lm EGDe in hydrogel, compared to planktonic cells. Moreover, bacteria organised in biofilms, contrary to a planktonic lifestyle, are known to be more tolerant and resistant to antibiotics^62^, and to the gastrointestinal tract^27,42^. To some extent, the divergent responses observed among Gram-positive and Gram-negative pathogens, highlight that a spatialised growth does not impose a single adaptive strategy, but rather interacts with species-specific physiology. That is why, we hypothesize that there is also metabolic regulation, extracellular polymeric substances production, stress response activation, and phenotypic heterogeneity in microcolonies, to promote survival under extreme conditions.

While *in vitro* assays revealed clear effects of spatialised growth against acidic stress, these differences were not reflected at the scale of the whole *H. illucens* larva by global enumeration (**Table 1**). Nevertheless, our results highlighted that at the scale of the acidic region of the dissected intestine, the same enhanced tolerance characteristics in spatialised lifestyle S-GSFM can be observed (**Figure 5**). This result demonstrates that the adaptive benefit conferred by spatialised growth is localised to the most selective digestive niche and can remain invisible when survival is evaluated at the whole-organism level. Moreover, some studies demonstrated a significant effect of biofilm spatialisation protection against gastrointestinal stresses, in a mouse model^27,63^, or in the nematode, *Caenorhabditis elegans*^64^. We demonstrate here that it is also the case for foodborne pathogens that are embedded in a gellified food-like matrix.

Our results also demonstrated that the presence of high osmotic pressure confers greater tolerance to *in vivo* digestive stresses, a property not observed in GSFM. In addition, *H. illucens* gut represents a highly heterogeneous environment, characterised not only by acidic pH, but also by digestive enzymes (amylase, lipase), antimicrobial components and a complex microbiota^65,66^. This suggests a cumulative effect of microcolonies biofilm and NaCl in protecting bacteria against all factors in the larval gut. A correlation can be done with the protective effect of a spatialised lifestyle against food stress and, more especially, osmotic stress. Mechanisms involved in osmotic stress responses may also be involved in digestive stress responses. Finally, this study reveals the potential of *H. illucens* larval as a model for gastrointestinal digestive simulation, with a specificity for acidic pH.

The findings of this study highlight the necessity to adopt approaches capable of capturing the spatial and physiological heterogeneity inherent to microcolonies. While conventional global population analyses provide important information, they mask localised responses that may confer adaptive advantages through spatialised growth. Recent methodological advances offer this opportunity^67^. Spatially resolved transcriptomics coupled with laser capture microdissection, allows precise sampling of cells from specific regions within biofilms, making it possible to directly link spatial position to gene expression patterns^68,69^. Other methodologies based on imaging, further improve our understanding of spatial mechanisms, by using Fluorescence In Situ Hybridisation (FISH)^51,70,71^. When combined with physiological or reporter-based probes, these approaches can reveal how metabolic activity, stress responses or growth states are distributed within a spatialised bacterial community ^52,72^. Finally, applied to food matrices, the integration of multi-modal approaches represents a particularly promising avenue.

Our findings challenge the implicit assumption that bacterial behaviour in foods can be reliably inferred from planktonic laboratory models. Spatial growth within structured matrices profoundly alters stress responses and digestive tolerance. Consequently, pathogen resilience may be systematically underestimated when spatial organisation and growth history are not considered. In quantitative microbial risk assessment, variability is commonly incorporated through strain diversity and host susceptibility, yet the physiological state of bacteria at ingestion remains largely overlooked. Our results indicate that spatial structuring in foods can alter digestive survival without necessarily affecting total population size, thereby potentially shifting the effective infectious dose.

Integrating spatial growth parameters into predictive microbiology frameworks therefore represents a critical step toward more realistic exposure assessment. Moving beyond homogeneous liquid paradigms toward structurally informed models may constitute a necessary evolution in predictive food microbiology.

## 4. MATERIALS AND METHODS

### 4.1. Bacterial strain and culture conditions

Eight major genera of foodborne bacterial pathogens relevant to food safety were selected to encompass a broad range of morphological and physiological characteristics. For some species, two strains were included to account for intraspecific variability. The complete strain list is presented in **Table 2**.

**Table 2:**
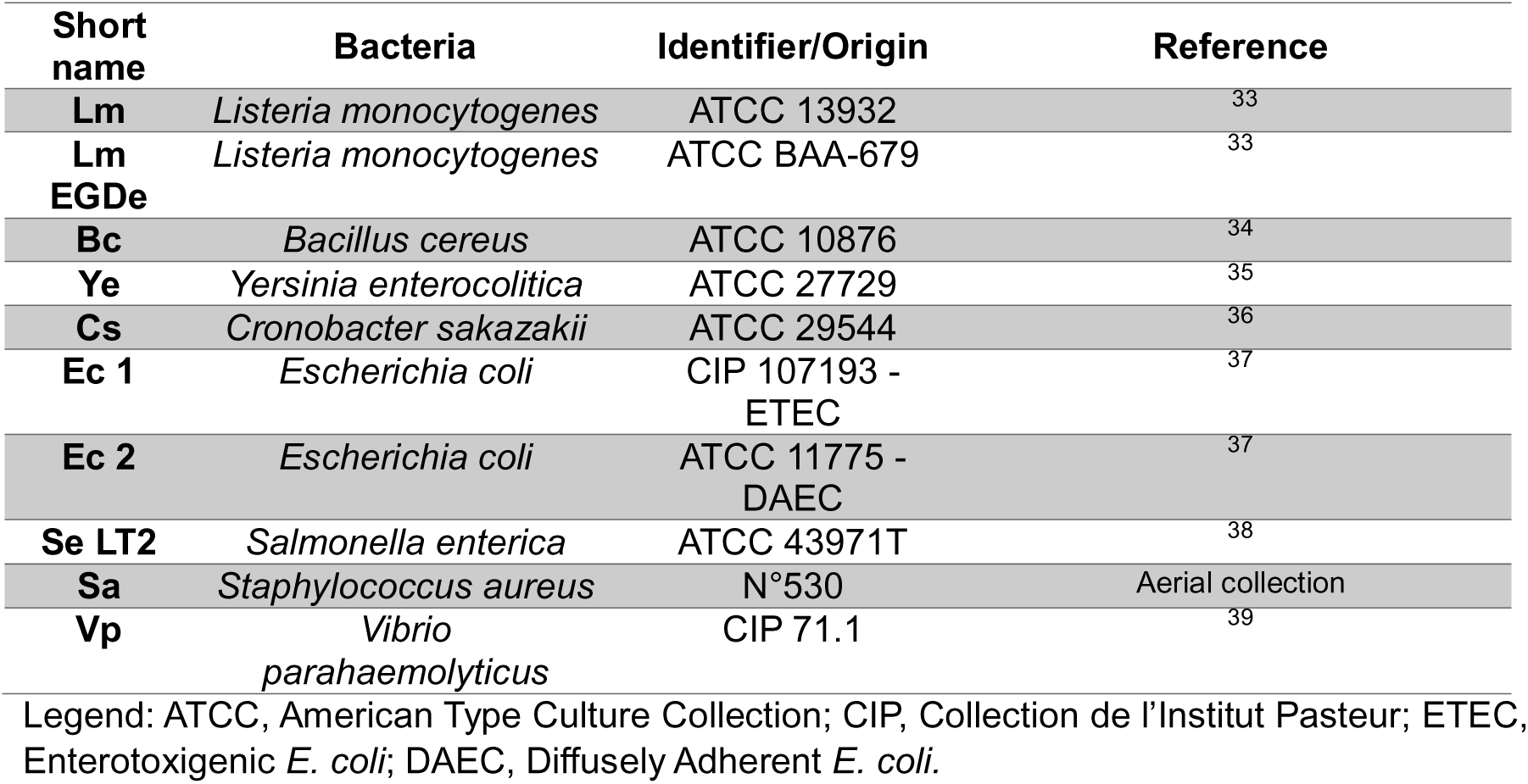
List of foodborne pathogens strains used in this study.

All strains used in this study were stored at -80°C in cryovial tubes containing glycerol media (Aerial collection). All cultures were performed in tryptone soy broth TSB 30 g/L (17.0 g/L of tryptic hydrolysate of casein; 3.0 g/L of soy peptone; 5.0 g/L of sodium chloride; 2.5 g/L of dipotassium phosphate and 2.5 g/L of glucose) (Themofisher, Dardilly, France) supplemented with 0.6% yeast extract (Thermofisher, Dardilly, France), TSBYe media, pH 7.3, sterilised by filtration using a 0.22 µm PET (Polyethene terephthalate) membrane (Thermofisher, Dardilly, France). Prior to each experiment, a culture was incubated at 30°C (Lm; Bc; Ye) or 37°C (Cs; Ec; Se; Sa; Vp) for 16 h. This culture was diluted at least twice at the centennial in TSBYe and incubated at the appropriate temperature (see above) for 8 h. This second culture was diluted to a minimum at the centennial, first in tryptone salt (Thermo Fisher Scientific, Dardilly, France) and then in TSBYe, and incubated at the appropriate temperature (see above) for 16 h.

The media described below (TSBYe) is used for planktonic lifestyle monitoring. Different media, called “stress medium”, were used in this study (**Table 3**).

**Table 3:** Percentages of ingredients/additives composing “stress medium”. The initial NaCl (Sigma-Aldrich, Saint Louis, Missouri, USA) concentration in the TSBYe medium was included in the desired final concentrations.

| Level | NaCl | Lactic acid | Acetic acid |
| --- | --- | --- | --- |
| 1 | 2.00 % | 0.25 % | 0.05 % |
| 2 | 3.00 % | 0.50 % | 0.10 % |
| 3 | 4.50 % | 0.80 % | 0.20 % |
| 4 | 6.00 % | 1.00 % | 0.50 % |
| 5 | 18.00 % | 2.00 % | 1.00 % |

For medium supplemented in lactic acid (Sigma-Aldrich, Saint-Louis, Missouri, USA) and acetic acid (Merck, Darmstadt, Germany), medium pH was adjusted to 7.4 with sodium hydroxide (NaOH) (Merck, Darmstadt, Germany) before sterilisation by filtration using a 0.22 µm polyethene terephthalate (PET) membrane (Thermofisher, Dardilly, France).

To assess the growth in spatialised lifestyle, adjusted “stress media” are used. They are prepared in a manner that after dilution to half in ultra-pure agarose (Invitrogen, France), sterilized by autoclaving, the final concentrations are 0.5% ultra-pure agarose; 30 g/L TSB; 0.6% yeast extract and the appropriate concentrations of ingredients/additives (see **Table 3**). Mixes were made at 40°C. The hydrogel also formed corresponds to a Gellified Synthetique Food Matrices (GSFM) containing 0.5% agarose. In the continuation of the study, the GSFM supplemented with 6.00% NaCl will be more extensively studied and named S-GSFM.

### 4.2. Growth monitoring protocols

#### 4.2.1. Planktonic cultures

Bacterial growth in a planktonic lifestyle was monitored using an automated turbidimetric system, Bioscreen C (Oy Growth Curves Ab Ltd., Raisio, Finland).

Bacterial cultures were recovered in the stationary phase and diluted to approximately 6 log_10_ CFU/mL in tryptone-salt medium. Three hundred microliters of “stress medium” was added to each well, and 30 µL of culture containing 6 log_10_ CFU/mL of bacteria was inoculated. The final concentration of bacteria in each well was approximately 5 log_10_ CFU/mL. The microplates were incubated in Bioscreen C at 25°C without shaking. Optical density (OD) measurements at 600 nm were taken every 10 minutes for a period long enough for all bacterial strains to reach the stationary phase. For the planktonic lifestyle, a total of 1,350 growth curves were acquired: 10 bacteria x 3 food-stress media x 5 levels x 3 technical replicates x 3 biological replicates (**Supplementary material 7**).

#### 4.2.2. Spatialised cultures

To account for bacterial spatial heterogeneity, multiple reading methods were developed to monitor the growth in hydrogels. Our temporal data on spatialised growths allow us to extract parameters similar to those of planktonic growth, enabling us to compare the two lifestyles.

Bacterial growth in a spatialised lifestyle was followed, in 96-wells CytoOne microplate (Starlab, Orsay, France) by using a Multiskan Ascent spectrometer (Thermofisher, Dardilly, France). Multipoint OD measurements (at 600 nm) were taken either at the central point of the well (model 1) or four points within the well (model 2). These four measurement points were precisely located by adjusting the coordinates of the plate: northeast, northwest, southeast and southwest of each well. Finally, we developed a multipoint reading technique that enables us to track the heterogeneity of spatialised growth.

The hydrogel model developed by Saint Martin et al., 2023, was used to follow pathogen growth in a spatialised lifestyle^26^. A 1:1 mix of each doubled-concentrated “stress medium” (500 µL) and ultra-pure agarose (500 µL) was prepared and stored at 40°C in an Eppendorf ThermoMixer® C (Eppendorf, Montesson, France) prior to inoculation, under agitation at 300 rpm. Bacterial cultures were recovered in the stationary phase (as described above) and diluted to approximately 6 log_10_ CFU/mL in tryptone-salt medium. Mixes (stress medium and agarose) were inoculated at 40°C with 100 µL of this dilution to obtain a final mix containing approximately 5 log_10_ CFU/mL of bacteria. Then, a 96-well CytoOne microplate was inoculated by pipetting 200 µL of the mix into each well, by taking care to avoid air bubbles in the agar. The surface of the agarose gel should be smooth and without slope in the well, so as not to induce a bias in growth tracking. Each mix (bacteria/stress medium pair) was done in triplicate, and three independent experiments were conducted (n = 9). Directly after inoculation, a first read with model 2 (4 points within the well) was performed to measure the initial OD. These OD measurements must be less than 0.094, as this indicates the presence of a slope or bubble in the well.

Then, the microplates are packed in a plastic bag to prevent drying and placed in an incubator at 25 °C under static conditions. Each hour, OD at 600 nm was manually read using the model 1, then the model 2. Growth was monitored over 3 consecutive days, and a final time point was collected after 6 days.

A total of 160 bacteria/stress medium pairs were analysed for their spatialised lifestyle, with 9 replicates. For each pair/well, a multipoint reading was performed, yielding 7,200 growth curves (**Supplementary material 8**). Finally, for each condition, the five measurement points are analysed to remove those corresponding to the development of a gas bubble or the absence of microcolonies. The curves for each well are averaged to generate a mean OD (ODm).

#### 4.2.3. Growth parameters extraction

Data processing and analyses were performed in R Version 4.3.2. Data processing was performed using the *Tidyverse* package, and growth curves were fitted using *Biogrowth*^40^. The Baranyi and Roberts model^41^, was used for estimating the Lag phase, Lag (h), and the specific maximum growth rate, *µ*_max_ (h^-1^). These parameters were used to characterise the bacterial growth and to estimate the lifestyle impact.

An automated R script was developed to extract the raw data, including the time in hours and the ODm. R codes for extract growth parameters are accessible *via* INRAE forge: https://forge.inrae.fr/elodie.hoch/code_availability_microcolonies/-/tree/34d0d59ba9e887c264e3c68a4b2f31ebfd04cd01/. First, to ensure a better fit of the model, the raw data are shortened to retain only about ten hours of ODm corresponding to the stationary phase of growth. Then, the initial parameters were estimated using the *make_guess_primary()* function and the model adjustment was performed with the *fit_growth()* function, with the environment parameter fixed to a *constant*. In the case where the initial adjustment failed or was unstable, the initial OD and maximal OD were fixed to the minimal OD and maximal OD, respectively, to stabilise the optimisation. All adjustments were visualised on the growth curves. Each adjustment parameter was saved in a final table. The whole results were exported to Excel format for a comparative analysis between planktonic et spatialised lifestyles (see **Supplementary material 1**).

#### 4.2.4. Statistics

The statistical analyses were performed using R version 4.3.2. Results are expressed as ΔLag and Δ*µ*_max_, corresponding to the difference between growth parameters in the control condition and in stress conditions. Statistics were performed on delta Lag and delta *µ*_max_ for each stress condition in the two lifestyles.

A R script was developed to automate the next steps. R codes for statistical analyses are accessible *via* INRAE forge: https://forge.inrae.fr/elodie.hoch/code_availability_microcolonies/-/tree/34d0d59ba9e887c264e3c68a4b2f31ebfd04cd01/. First, the possibility of applying a parametric test is assessed by checking the normality of the data using the Shapiro-Wilk test and the homogeneity of variances using Levene’s test. If the variances are equal and the data follow a normal distribution, a Student *t*-test is used. Conversely, if the variances are unequal but the data still follow a normal distribution, Welch’s *t*-test is used. Finally, if the samples do not follow a normal distribution, the non-parametric Mann–Whitney test is applied. An Excel file was generated, regrouping the p-value for each test, for each strain and stress conditions. An appropriate test was selected for each sample, and the p-values were indicated by asterisks on the graph.

### 4.3. Digestive stress impact assessment

The following culture preparation protocol is applicable to both *in vitro* and *in vivo* models. Bacterial cultures of Lm EGDe, Bc, Ye, and Se LT2 were recovered in the stationary phase (as described previously) and diluted to approximately 6 log_10_ CFU/mL in tryptone salt. In this part, three hydrogels were tested: GSFM, GSFM 2.00% NaCl and GSFM 6.00% NaCl (S-GSFM). To evaluate the impact of digestive stress on bacteria living in a planktonic lifestyle, 1 mL of the suspension was used to inoculate 9 mL of the previously mentioned medium. To compare the impact of digestive stress on bacteria with a spatialised lifestyle, cultures were prepared in the same medium as previously described in Section 2.2.2. All samples were incubated at 25°C under static conditions for 96 hours.

#### 4.3.1. *In vitro* digestive stress model

The following protocol was used to evaluate the impact of *in vitro* digestive stress. All steps were applied for each strain tested. For a spatialised lifestyle, wells were collected individually into 2 mL Eppendorf tubes or 15 mL Falcon tubes. For planktonic cultures, 200 µL of culture was collected from each strain into 2 mL Eppendorf tubes or 15 mL Falcon tubes. Planktonic cultures were centrifuged at 5,000 x g for 5 min at 4°C. Supernatants were removed, and the pellets were resuspended in digestion solution (stress or control), or in spatialised cultures (the well was collected). Digestive stress solutions were prepared as described by Chamignon et al., 2020^42^. All solutions were filter-sterilised by using a 0.22 µm PET (Polyethene terephthalate) membrane. The following manipulation plan was followed for each bacterial strain, in biological triplicate:

1. Sample 1: 1.8 mL of 1% bile salts solution (9 g/L NaCl, 10 g/L bile salts (Sigma-Aldrich, Saint-Louis, Missouri, USA)). Incubate 2 h at 37°C, under agitation at 160 rpm.
2. Sample 2: 1.8 mL of 10 mM H_2_O_2_ solution (9 g/L NaCl, 0,34 g/L H_2_O_2_ (Thermofisher, Dardilly, France)). Incubate 2 h at 37°C, under agitation at 160 rpm.
3. Sample 3: 27 mL of physiological solution (9 g/L NaCl), adjusted at pH 2.0 with hydrogen chloride (HCl) 5 M (Sigma-Aldrich, Saint-Louis, Missouri, USA). Incubate 4 h at 37°C, under agitation at 160 rpm.
4. Sample 4: 1.8 mL of physiological water solution (9 g/L NaCl), adjusted at pH 7.0 with HCl 5 M. Incubate 2 h at 37°C, under agitation at 160 rpm.
5. Sample 5: 27 mL of physiological water solution (9 g/L NaCl), adjusted at pH 7.0 with HCl 5 M. Incubate 2 h at 37°C, under agitation at 160 rpm.

After incubation, all samples were centrifuged at 5,000 x g for 5 min at 4°C. Supernatants were removed, and the pellets were resuspended in 200 µL saline solution (9 g/L NaCl). Samples corresponding to planktonic lifestyle are directly inoculated on TSYa (17.0 g/L of tryptic hydrolysate of casein; 3.0 g/L of soy peptone; 5.0 g/L of sodium chloride; 2.5 g/L dipotassium phosphate; 2.5 g/L of glucose medium and 15 g/L agar), by spot method (10 µL), by performing 10-fold serial dilutions from 10^-2^ to 10^-8^.

For the sample corresponding to a spatialised lifestyle, wells were broken mechanically by using a loop. The samples were vortexed to release cells from the agar matrix and resuspend them in physiological saline. One millilitre of physiological saline was added to each sample, and the mixture was filtered in a new Eppendorf tube using a cell strainer with a 20 µm porosity to remove agar debris. Suspensions collected are centrifuged at 5,000 x g for 5 min at 4°C. Supernatants were removed, and pellets were resuspended in 200 µL physiological water. Samples were then inoculated into TSYa using the same protocol as previously described for the planktonic sample. Petri dishes were incubated for 48 h at 30°C.

The impact of digestive stress was assessed by measuring cultivability loss. The following equations were applied:

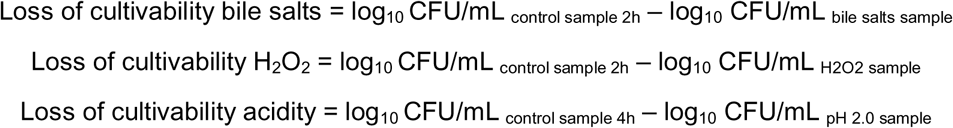

#### 4.3.2. *In vivo* digestive stress model

A larval model, the black soldier fly *Hermetia illucens*, was selected to evaluate the *in vivo* impact of digestive stress, specifically the acidity. Indeed, *H. illucens* has a digestive tract in which pH varies greatly depending on the region of the intestine^43^. An ad libitum ingestion model was developed to assess the *in vivo* impact of this digestive stress, as described below. Due to the complexity of the *in vivo* model, experiments were restricted to *Salmonella enterica* serovar Typhimurium (Se LT2), as a model organism.

This study utilised larvae reared at Institut de Recherche sur la Biologie de l’Insecte, UMR7261 CNRS, Université de Tours (Tours, France) and delivered to GME Micalis institute INRAE laboratory at Jouy-en-Josas at the juvenile stage (2 - 5 mg), housed in a container containing a mix of ground wheat, corn and lucerne at 70% humidity, allowing further development prior to experimentation. For the *ad libitum* ingestion model, larvae weighing 30-45 mg each were selected. Experiments were conducted at 25°C in 90-mm-diameter Petri dishes sealed with Parafilm.

An additional preparation step, beyond what has been previously described, was necessary for *in vivo* model. Previously prepared planktonic cultures (TSBYe, TSBYe 6.00% NaCl) were collected and centrifuged at 5,000 x g for 5 min at 4°C. Supernatant was then recovered and filtered through a 0.22 µm membrane. Three mixes were prepared, each containing 500 µL of medium, 500 µL of 1% agarose, and 100 µL of physiological water. Subsequently, 200 µL of each mix was inoculated in triplicate into a 96-well plate. Samples were incubated at 25°C under static conditions for 96 hours.

For planktonic lifestyle, four-day cultures were centrifuged at 5,000 x g for 5min at 4°C, and pellets were resuspended in an equal volume of physiological saline. According to the condition, one uninoculated well is recovered and placed in a 90-mm-diameter Petri dish. Thirty-three microliters of cell suspension were dispersed on the well.

For a spatialised lifestyle, one inoculated well is recovered and placed in a 90-mm-diameter Petri dish. Larvae were disinfected in 70% ethanol, then in sterile physiological saline, before being added to one larva per Petri dish. Finally, the ratio of larvae/feed (well of medium) was 1:1. Ad libitum ingestion was allowed for 24 h. Controls were added to this experiment to verify the absence of native *Salmonella* in *H. illucens* (control 1), to measure the initial level of Se LT2 added as food (control 2) and the final level of Se LT2 without larvae (after 24h at 25°C) (control 3). The experiment was performed four times, corresponding to four biological replicates.

After 24 hours, larvae were recovered and subjected to a 30-minute cooling period. The larval disinfection process was performed to remove the pathogens from the cuticle (larval skin). In a first experiment, three larvae per condition were crushed in 300 µL cold physiological saline, and *S. enterica* counts (CFUs) were determined through serial 10-fold dilution and plating on CHROMagar^TM^ Salmonella medium (Saint-Denis, France), which enabled the detection of *S. enterica,* appearing as purple colonies. After removing the larvae, the Petri dish was washed with 1 mL physiological saline to collect the Se LT2 that had not been eaten. Suspension was plated on CHROMagar^TM^ Salmonella medium by serial 10-fold dilutions. Plates were incubated at 37°C for 24 hours. In a second experiment, three larvae per condition were independently dissected, using sterile tweezers and scissors to avoid cross-contamination of the samples. Each gut was isolated in sterile physiological saline water in a sterile 90 mm diameter Petri dish. Once collected, the gut was divided into three regions, anterior, middle and posterior. Only the acid (pH 2) middle gut region was collected, with a sterile scalpel and placed into 300 µL 30% glycerol physiological saline water, until plating. The gut was disrupted by pipetting up and down and suspension was plated on CHROMagar^TM^ Salmonella medium by serial 10-fold dilutions.

#### 4.3.3. Statistics

The statistical analyses were performed using XLSTAT version 2025.1. Results are expressed as the mean ± the standard deviation error of three (*in vitro*) or four (*in vivo*) independent experiences. For *in vitro* essays, an analysis of variance (ANOVA) test was performed to determine whether there were any significant differences among the means of loss of cultivability, at a 95.0% confidence level (α = 0.05). If the ANOVA indicated significant differences in the impact of digestive stress between the two lifestyles, a Fisher least significant difference (LSD) test was used to identify which means were significantly different. For *in vivo* essays, a non-parametric Mann-Whitney test was performed to determine whether there were any significant differences between the two lifestyles and the presence of a high concentration of NaCl.

### 4.4. Confocal Laser Scanning Microscopy (CLSM)

All microscopic observations were performed with a Leica HCS-SP8 confocal laser scanning microscope at the INRAE MIMA2 imaging platform (https://mima2.jouy.hub.inrae.fr/). For observations, strains were tagged fluorescently in red with SYTO 61 (1:500 dilution in TSBYe medium from a stock solution at 5 µM in DMSO (Invitrogen, France)), a nucleic acid dye. SYTO 61 excitation was performed at 633 nm using an HeNe laser, and the emitted fluorescence was recorded within the ranges of 640-690 nm a photomultiplier detector. To capture the morphology and repartition of microcolonies, a 20x/0.75 DRY HC PL APO CS2 objective was used with a 512 x 512-pixel definition. A bidirectional acquisition speed of 600 Hz allowed a frame rate of 2.3 images per second. For 3D stack analyses, a 1µm step was used between z levels. For each condition, 3 stacks per well were acquired with a height of 300 µm. Image analysis was processed using IMARIS 9.3.1 (Bitplane, Switzerland).

Background fluorescence was first subtracted from all pictures before applying a Gaussian filter to the red fluorescence. Raw fluorescence of each image was then binarised using the automated Imaris isosurface function. Geometric parameters (e.g. biovolume, sphericity, surface area) were extracted from these binarised images.

### 4.5. Automated temporal imaging of microcolonies

For macroscopic observations of microcolonies formed in hydrogel medium, 12-well plates were placed into the Reshape Biotech ImageRead system (Reshape ApS, Denmark). It is an automated platform that integrates robotic incubation, high-resolution imaging, and real-time, AI-powered, deep-learning quantitative analysis. The equipment is specifically designed for detecting low-density, small-sized colonies. Small colonies were modelled using the SBS-format 12-well plates for this study. Imaging was carried out under controlled incubation at 25°C, using a combination of top and bottom lighting to enhance optical contrast. The imaging system acquired one image per hour automatically for five days (120 hours).

## 5. CONCLUSION

Spatialised growth in food-like matrices fundamentally reshapes the physiological state of foodborne pathogens in a species- and stress-dependent manner. By promoting microcolony formation, spatial constraints enhance resilience to both food-related and digestive stresses compared with planktonic cultures. Crucially, this adaptive advantage may remain undetected when survival is assessed at a bulk scale. Extending principles established in biofilm biology to food-embedded microcolonies provides a mechanistic bridge between food microstructure, bacterial growth history and infection risk. Recognising spatial organisation as a determinant of pathogen fate highlights the need to incorporate structural context into food safety studies and risk assessment frameworks.

## Acknowledgements

This research was funded by AERIAL, INRAE, and “Association Nationale de la Recherche et de la Technologie” (contract 2023/0850). We thank the MIMA2 imaging platform (Microscopie et Imagerie des Microorganismes, Animaux et Aliments, https://doi.org/10.15454/1.5572348210007727E12) for the Leica SP8-HCS microscopy observations. Figure 3B was created with BioRender (https://biorender.com). Adrienne Lintz and Sarah Côte are acknowledged for technical assistance. Carlotta Savio is acknowledged for her help in developing the insect larval model. Cecile Berdous for technical assistance on larval dissections.

## Author contributions

E.H., L.F.O. and R.B. conceptualised the overarching aims of the research study. E.H., B.H., L.F.O. and R.B. conceived and designed the experiments. E.H., C.N.L., performed the experiments and data acquisition. E.H., C.N.L., L.G., B.H., L.F.O. and R.B. analysed and interpreted the data. L.F.O. and R.B. had both management and coordination responsibilities for the execution of the research work. L.F.O. and R.B. contributed to the acquisition of the financial support and resources leading to this publication. All authors substantially (i) contributed to the conception or design of the work or the acquisition, analysis or interpretation of the data, (ii) revising critically the manuscript for important intellectual content and (iii) approved the final approval of the completed version.

