## Supplementary Material for "Spatial growth in food-like matrices differentially modulates food-related stress responses but enhances digestive tolerance in major foodborne pathogens"

### Supplementary materials

**Supplementary table 1:** Lag and μmax values extracted from Barany's and Roberts model for each bacteria/stress pairs (n=9).

**Supplementary table 2**: Table showing the distribution of strains (**A**) and food-related stresses (**B**) across the 3 clusters of the K-means clustering of microbial growth parameters.

**Supplementary material 3:** Reshape videos of pathogens microcolonies growing in GSFM and S-GSFM during 4 days at 25°C.

**Movie S3a:** Growth of Se LT2 microcolonies in GSFM medium during 4 days at 25°C. Video monitored by Reshape Biotech. One image was taken each hour in a 22.6 mm Ø well (Final video accelerated 4x, final time 3s).

**Movie S3b**: Growth of Se LT2 microcolonies in S-GSFM medium during 4 days at 25°C. Video monitored by Reshape Biotech. One image was taken each hour in a 22.6 mm Ø well (Final video accelerated 4x, final time 3s).

**Movie S3c**: Growth of Ye microcolonies in GSFM medium during 4 days at 25°C. Video monitored by Reshape Biotech. One image was taken each hour in a 22.6 mm Ø well (Final video accelerated 4x, final time 3s).

**Movie S3d**: Growth of Ye microcolonies in S-GSFM medium during 4 days at 25°C. Video monitored by Reshape Biotech. One image was taken each hour in a 22.6 mm Ø well (Final video accelerated 4x, final time 3s).

**Movie S3e**: Growth of Lm EGDe microcolonies in GSFM medium during 4 days at 25°C. Video monitored by Reshape Biotech. One image was taken each hour in a 22.6 mm Ø well (Final video accelerated 4x, final time 3s).

**Movie S3f**: Growth of Lm EGDe microcolonies in S-GSFM medium during 4 days at 25°C. Video monitored by Reshape Biotech. One image was taken each hour in a 22.6 mm Ø well (Final video accelerated 4x, final time 3s).

**Movie S3g**: Growth of Bc microcolonies in GSFM medium during 4 days at 25°C. Video monitored by Reshape Biotech. One image was taken each hour in a 22.6 mm Ø well (Final video accelerated 4x, final time 3s).

**Movie S3h**: Growth of Bc microcolonies in S-GSFM medium during 4 days at 25°C. Video monitored by Reshape Biotech. One image was taken each hour in a 22.6 mm Ø well (Final video accelerated 4x, final time 3s).

**Supplementary material 4**: Dissection of the intestinal gut of *H. illucens* after 24 hours ad libitum ingestion of GSFM containing Se LT2 (Petri dish ø 60 mm).

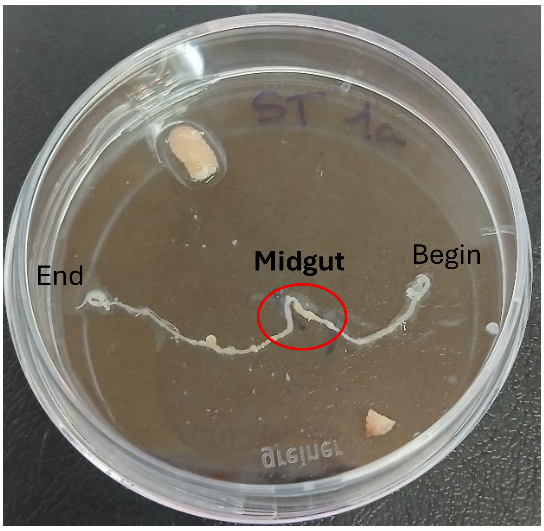


**Supplementary material 5**: pH measurements of the different regions (Foregut, midgut, and hindgut) of *H. illucens* intestinal gut.

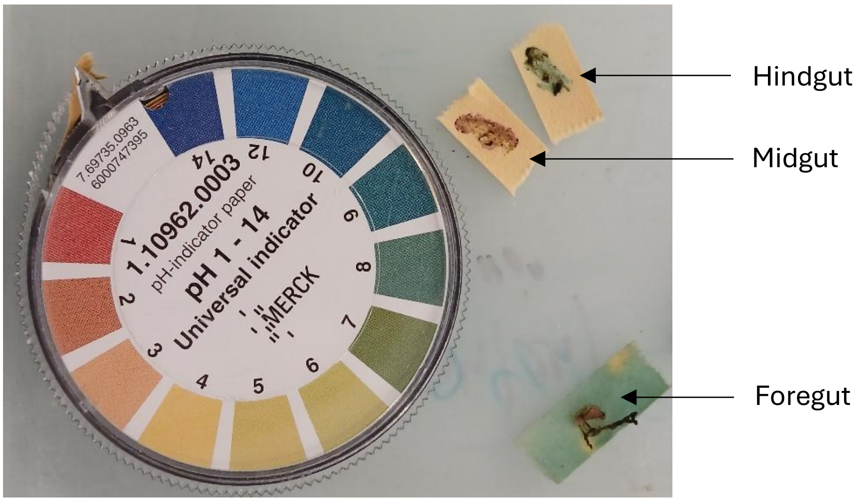


**Supplementary material 6**: Videos of *H. illucens* ad libitum ingestion model.

**Movie S6a**: Ingestion of planktonic cells of Se LT2 in GSFM by *H. illucens* in an ad libitum ingestion model. One image was taken each ten minutes in a 90 mm Ø well (Final time 10s).

**Movie S6b**: Ingestion of spatialised cells of Se LT2 in GSFM by *H. illucens* in an ad libitum ingestion model. One image was taken each ten minutes in a 90 mm Ø well (Final time 10s).

**Movie S6c**: Ingestion of GSFM control without bacteria in *H. illucens* in an ad libitum ingestion model. One image was taken each ten minutes in a 90 mm Ø well (Final time 10s).

**Supplementary table 7:** Growth curves obtained in planktonic lifestyles for each bacterial stress pairs OD=f(time) (n = 9). Each workbook corresponds to a strain. The durations of growth monitoring are not homogeneous between replicates; they are adapted according to the speed of growth.

**Supplementary table 8:** Growth curves obtained in spatialised lifestyles for each bacterial stress pairs ODm=f(time) (n = 9). Each workbook corresponds to a strain. The durations of growth monitoring are not homogeneous between replicates; they are adapted according to the speed of growth.
